# Like host, like parasite: intraspecific divergence in a polystomatid flatworm parasite across South Africa echoes that of its frog host

**DOI:** 10.1101/2022.03.15.483565

**Authors:** Anneke Lincoln Schoeman, Nikol Kmentová, Maarten PM Vanhove, Louis Heyns Du Preez

## Abstract

The African Clawed Frog *Xenopus laevis*, a global invader, exhibits marked phylogeographic divergence among native populations in southern Africa, which enhances its invasive potential. The polystomatid flatworm *Protopolystoma xenopodis*, as the frog’s most frequently co-introduced metazoan parasite, may be the ideal biological tag for the frog’s movement, if corresponding divergence can be demonstrated. In an integrative approach, we utilised morphometrics and molecular markers to assess divergence in *P. xenopodis* in its native range. We measured twelve key morphological characters from 23 flatworms and compared these statistically between flatworms collected to the north and south of the Great Escarpment Mountain Range in South Africa. Phylogenetic analyses were based on three concatenated markers, namely *28S* and *12S rDNA* and *COX1*, from six flatworms. The combination of five morphological characters, which involve egg size, gut morphology and size of the attachment hooks, differentiated northern and southern populations of *P. xenopodis* in South Africa. The multilocus phylogenetic analyses supported these findings, showing a well-supported cluster of northern *P. xenopodis*. These findings suggest that taxonomic studies of polystomatid flatworms should make use of geographically representative data sets that consider both morphological and molecular evidence. Moreover, the findings demonstrate that the frog host and flatworm parasite exhibit corresponding phylogeographic structuring in the native range. Consequently, the phylogeography of *P. xenopodis*, both in the native and invasive range of its host, may act as a key piece of evidence to reconstruct past invasion pathways of *X. laevis*.

## 1 INTRODUCTION

Variability, which promotes the adaptability and viability of populations in changing environments, is a factor to be reckoned with when it comes to the invasion success of alien species, as has been shown for several taxonomic groups in aquatic ecosystems (Wellband *et al*., 2017). Consequently, a thorough understanding of the evolutionary history of an invasive species in its native range is essential to assess its potential to colonise and adapt to novel surroundings (Lee & Gelembiuk, 2008). This is equally true for the co-introduced parasites of free-living invasive species, which make out most non-native species (Torchin *et al*., 2003). Yet, alien parasites are often overlooked in the study of biological invasions (Blackburn & Ewen, 2017).

Worldwide, the African Clawed Frog *Xenopus laevis* (Daudin 1802) (Anura: Pipidae) is one of the most widespread amphibians. Its native range covers much of sub-Saharan Africa (Furman *et al*., 2015). Invasive populations of this frog can be found in Asia, Europe and North and South America (Measey *et al*., 2012). Wherever *X. laevis* occurs, it harbours a diverse and unique parasite fauna (Tinsley, 1996). One of *X. laevis*’ most prevalent parasites is the host-specific flatworm *Protopolystoma xenopodis* (Price, 1943) (Monogenea: Polystomatidae), a sanguinivorous inhabitant of the frog’s bladder in its adult form. In the native range of southern Africa, *P. xenopodis* is a common feature of *X. laevis* parasite assemblages, where it has been recovered from more than 90% of *X. laevis* populations and more than 50% of all sampled hosts in a recent survey (Schoeman *et al*., 2019). Moreover, in the context of the global invasive status of *X. laevis, P. xenopodis* emerges as its most frequently co-introduced metazoan parasite and has been reported from hosts in France, Portugal, the United Kingdom and the United States (Tinsley & Jackson, 1998b; Kuperman *et al*., 2004; Rodrigues, 2014; Schoeman *et al*., 2019).

In general, *P. xenopodis* is differentiated from its congeners, which infect other *Xenopus* species in Africa, based upon the morphology of the gut, large posterior attachment hooks and spines on the male reproductive organs (Tinsley & Jackson, 1998b). The gut of *Protopolystoma* spec. bifurcates after the pharynx into two caeca, which branch out even further into diverticula that may fuse to form post-ovarian inter-caecal anastomoses (Tinsley & Jackson, 1998b). The number of diverticula and anastomoses varies within and between species (Tinsley & Jackson, 1998b). All *Protopolystoma* spec. possess a pair of large hooks, or hamuli, with two roots and a sharpened terminal hook, used to attach to the wall of the host’s bladder (Tinsley & Jackson, 1998b). Due to their size, complex shape and sclerotisation, the morphology of the large hamuli is the most taxonomically informative feature among members of the genus (Tinsley & Jackson, 1998b). Finally, *Protopolystoma* spec. have a muscular, bulb-shaped male reproductive organ armed with sixteen spines that are arranged in two concentric rings of eight spines each (Tinsley & Jackson, 1998b). It is hard to obtain reliable measurements of these spines but *P. xenopodis* appears to have much shorter spines than its congeners (Tinsley & Jackson, 1998b). In their redescription of *P. xenopodis*, Tinsley & Jackson (1998b) noted geographical variation in the size of the genital spines between southern and more northerly populations across sub-Saharan Africa. The authors also noted marked intraspecific variation in the morphology of the large hamulus and the caecal branches, although this was not correlated with geographic distance (Tinsley & Jackson, 1998b).

Given the wide distribution of this parasite across the globe, possible cryptic diversity, explored through both morphological and molecular data, is worth investigating as an essential building block of bio-invasion research (Mazzamuto *et al*., 2016). In addition to illuminating the evolutionary potential of this alien parasite, the exploration of the intraspecific divergence in *P. xenopodis* is worthwhile in the light of the phylogeographic structuring of its host *X. laevis* in its native range (de Busschere *et al*., 2016; Furman *et al*., 2015). Previous studies on the morphology and genetics of *X. laevis* have identified marked divergence among populations in southern Africa (Grohovaz *et al*., 1996; Measey & Channing, 2003; du Preez *et al*., 2009; Furman *et al*., 2015; de Busschere *et al*., 2016).

The intimate association between frog host and flatworm parasite would lead us to expect corresponding morphological and phylogenetic divergence among the populations of *P. xenopodis* across southern Africa. Congruence in host-parasite phylogeographies arises as a result of high host-specificity and direct life cycles, in combination with limited host-independent dispersal capacity (Nieberding & Olivieri, 2007). What is more, since parasites exhibit shorter generation times and greater abundance than their hosts, their phylogeographic divergence is often more pronounced (Nieberding & Olivieri, 2007). Consequently, parasites can act as biological magnifying glasses to further explore the host’s phylogeographic structuring and movement in both its native and invasive ranges (Nieberding *et al*., 2004). For example, the intraspecific morphological variation of monogenean flatworm parasites provided information on the invasion history of fish in Africa and Europe (Kmentová *et al*., 2019; Ondračková *et al*., 2012). In this framework, widespread co-introduced parasites of invasive hosts, such as *P. xenopodis*, could be ideal tags to trace the translocation of host lineages—if it can be demonstrated that they diverge according to a similar pattern as their hosts.

Therefore, the present study offers an exploratory investigation of the morphological differences and phylogenetic divergence in *P. xenopodis* collected from *X. laevis* from the northernmost or southernmost northern-most and southernmost localities in South Africa, linked to two distinct phylogeographic lineages according to de Busschere *et al*. (2016). In an integrative approach, we will rely on a combination of evidence from one nuclear and two mitochondrial genes and twelve key morphological characters to assess differentiation in *P. xenopodis* between the two regions. We expect (1) marked intraspecific variability in *P. xenopodis* in South Africa, (2) with significant divergence between northern and southern parasites in some taxonomically important morphological characters, such as gut morphology and dimensions of the sclerites, and in the three molecular markers, *COX1* and *12S rDNA* and *28S rDNA,* (3) which corresponds to the divergence of the host *X. laevis*.

## 2 METHODS

### 2.1 Specimen collection

From March to July 2017, 20 adult *X. laevis* were captured in funnel traps baited with chicken liver at eight field sites across South Africa (Table 1). These sites were located near previously sampled localities where the local *X. laevis* populations were genetically identified as belonging to either one of two phylogeographic lineages of this frog by de Busschere *et al*. (2016), namely SA1 to the southwest and SA5 to the northeast of southern Africa (Figure 1). These two groups of sites lie on either side the Great Escarpment, the edge of an inland plateau that runs continuously along the southwestern seaboard of southern Africa, which has been suggested as a natural barrier to gene flow for *X. laevis* (Furman *et al*., 2015). Based upon previous phylogeographic work, we can expect distinct lineages of this frog to the southwest, a winter rainfall region, and the northeast, a summer rainfall region, of the Escarpment (Furman *et al*., 2015).

**FIGURE 1.**
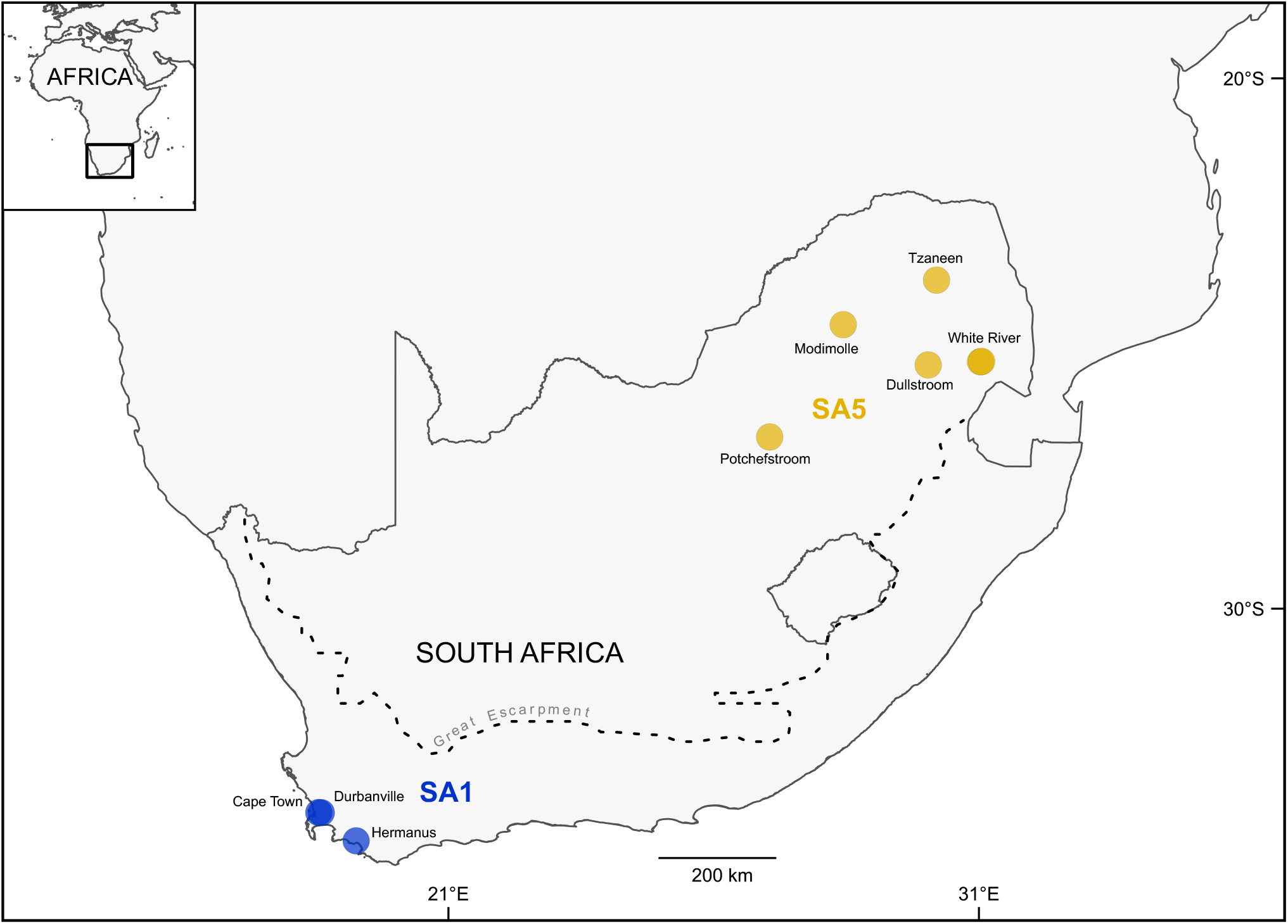
The eight localities where *Protopolystoma xenopodis* was obtained from *Xenopus laevis*, coloured according to the frogs’ expected phylogeographic lineage according to de Busschere *et al*. (2016), namely SA1 (in blue) to the southwest of the Great Escarpment Mountain Range (dotted line) or SA5 (in yellow) to the northeast. The locality names, derived from the nearest town, are indicated. The map was constructed in QGIS *version* 3.10.2-A Coruña (QGIS Development Team, 2018) with the Mercator projection.

**TABLE 1.** Information on the geographic origin of *Xenopus laevis* and their associated *Protopolystoma xenopodis* specimens. Localities are assigned according to the expected phylogeographic lineage of *X. laevis* in South Africa (de Busschere *et al*., 2016). Locality names refer to the nearest town and are given along with the collection date, geographic coordinates of the sampled water bodies, number of adult *X. laevis* hosts captured (N_X_) and number of *P. xenopodis* parasites collected for morphometry (N_P[m]_) and DNA sequencing (N_P[s])_.

| Locality | Date | Coordinates | $N_X$ | $N_{P[m]}$ | $N_{P[s]}$ |
| --- | --- | --- | --- | --- | --- |
| **SA1 host lineage (southwestern South Africa)** |  |  |  |  |  |
| Cape Town, Western Cape | June 2017 | 33.8355°S; 18.5528°E | 2 | 2 | 1 |
| Durbanville, Western Cape | July 2017 | 33.8392°S; 18.6003°E | 2 | 4 | 0 |
| Hermanus, Western Cape | June 2017 | 34.3702°S; 19.2571°E | 3 | 4 | 1 |
| **SA5 host lineage (northeastern South Africa)** |  |  |  |  |  |
| Dullstroom, Mpumalanga Province | April 2017 | 25.3981°S; 30.0380°E | 4 | 3 | 1 |
| Modimolle, Limpopo Province | April 2017 | 24.6384°S; 28.4369°E | 2 | 2 | 1 |
| Potchefstroom, North-West Province | March 2017 | 26.7555°S; 27.0506°E | 3 | 3 | 1 |
| Tzaneen, Limpopo Province | May 2017 | 23.7988°S; 30.1951°E | 2 | 2 | 1 |
| White River, Mpumalanga Province | April 2017 | 25.3391°S; 31.0226°E | 1 | 1 | 0 |
|  |  | 25.3320°S; 31.0433°E | 2 | 2 | 0 |

The frogs underwent double euthanasia according to institutional ethics guidelines under ethics approval number NWU-00380-16-A5-01: first anaesthesia in 6% ethyl-3-aminobenzoate methansulfonate (MS222) (Sigma-Aldrich Co., USA) and then euthanasia through pithing.

Frogs were dissected and adult specimens of *P. xenopodis* were obtained from the excretory bladder. The 29 retrieved polystomatids were processed for either morphological or molecular analyses (Table 1).

### 2.2 Morphometrical analyses

In total, 23 of the retrieved polystomatids from the eight localities were processed for morphological analyses. The live polystomatids were placed in a drop of tap water on a microscope slide and gently heated from underneath until they relaxed, following Snyder & Clopton (2005). They were then fixed in 10% neutral buffered formalin or 70% ethanol under coverslip pressure. Polystomatids preserved in both 10% neutral buffered formalin and 70% ethanol were hydrated through a decreasing ethanol series to tap water, with 10 minutes spent on each step. The specimens were stained overnight in acetocarmine. Thereafter, the specimens were dehydrated in an increasing ethanol series to absolute ethanol, 10 minutes per step, with colour corrections by hydrochloric acid incorporated whilst the specimens were in the 70% ethanol. The specimens were cleared in xylene and mounted in Canada balsam (Sigma-Aldrich Co., Steinheim, Germany). The mounts were dried at 50°C for approximately 48 hours.

Measurements and photomicrographs were taken on a Nikon ECLIPSE E800 compound microscope in conjunction with the software NIS-Elements Documentation *version* 3.22.09 (Nikon Instruments Inc., Tokyo, Japan). The following nine characters were measured: body length from the tip of the haptor to tip of the false oral sucker, body width at the widest point, length and width of the haptor, length of the ventral roots of the two large hamuli, length of the dorsal roots of the two large hamuli, length of the terminal hooks of the two large hamuli and the length and width of the egg (if present) at the longest and widest points, respectively (Figure 2). The following three structures were counted: number of post-ovarian inter-caecal anastomoses, number of medial diverticula of the caecum and number of lateral diverticula (Figure 2). The hamuli and the medial and lateral diverticula from the two sides of the polystomatids were measured or counted separately and then averaged for each specimen to give a single value for each character for subsequent analyses.

**FIGURE 2.**
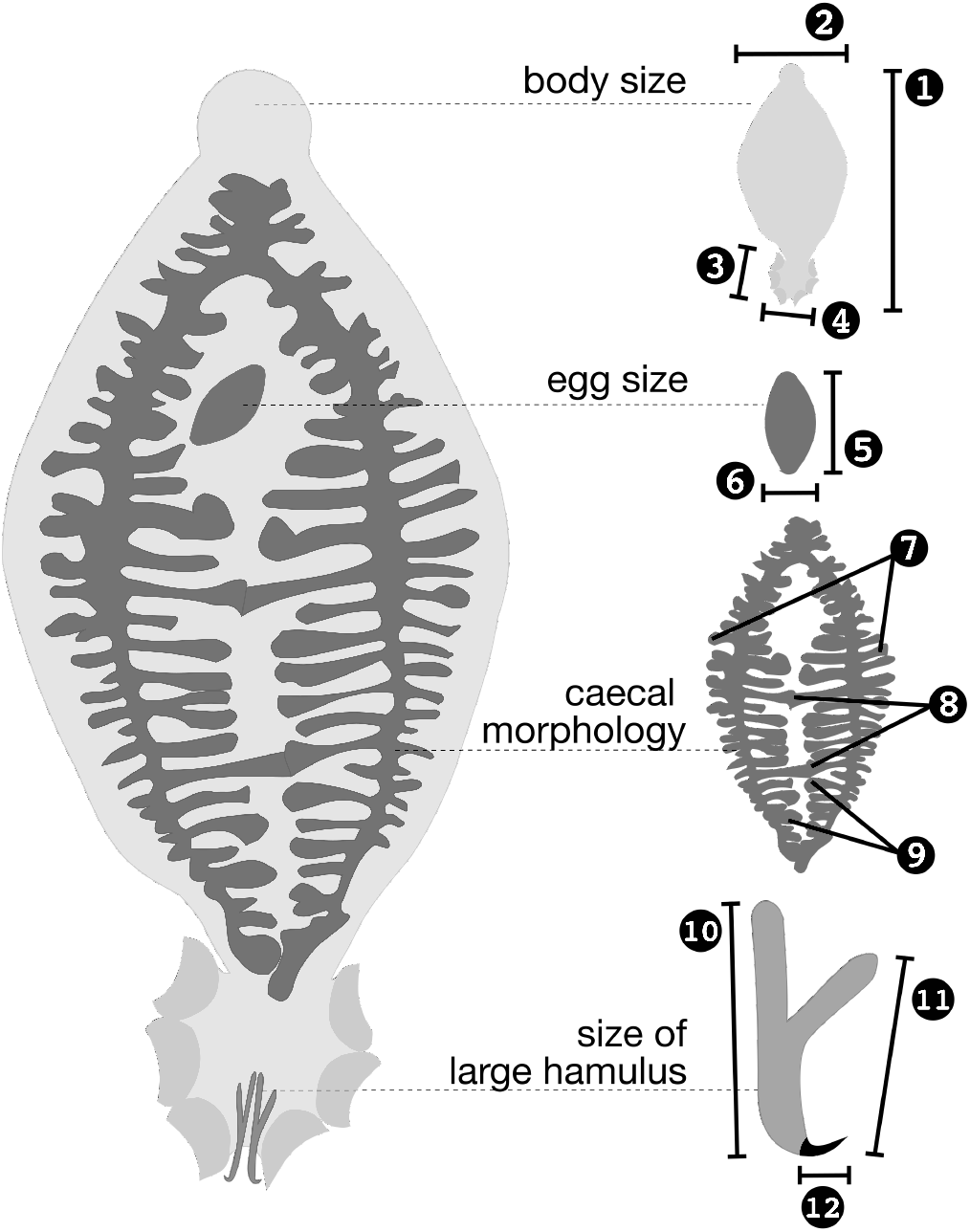
Measured morphological characters of adult *Protopolystoma xenopodis:* (1) body length, (2) body width, (3) haptor length, (4) haptor width, (5) egg length, (6) egg width, (7) number of lateral diverticula, (8) number of post-ovarian inter-caecal anastomoses, (9) number of medial diverticula, (10) length of the dorsal root of the large hamulus, (11) length of the ventral root of the large hamulus, (12) length of the terminal hook of the large hamulus.

The 12 characters were compared statistically based upon geographic origin (SA1 or SA5) in the software R *version* 4.1.2 (R Core Team, 2021). Unless otherwise mentioned, data carpentry and visualisation were performed with the help of the R packages *broom* (Robinson *et al*., 2022), *factoextra* (Kassambara & Mundt, 2020), *ggdist* (Kay, 2021), *ggtext* (Wilke, 2020), *patchwork* (Lin Pedersen, 2020), *png* (Urbanek, 2013), *skimr* (Waring *et al*., 2021) and *tidyverse* (Wickham *et al*., 2019). Missing data points were imputed by the random forest method in the R package *missForest* (Stekhoven, 2013) using a random seed of 666 as starting point. This method was preferred since it has a non-parametric approach suitable to the small sample size and because it can handle mixed variable types (Stekhoven & Bühlmann, 2012). Since certain characters can vary with parasite age, or its proxy, body size, the median body length and width and haptor length and width were compared between the two groups with the non-parametric Wilcoxon-Mann-Whitney (WMW) test to ensure that the groups contained polystomatids of similar size distributions. The WMW test was further employed to test whether there was a significant difference in the median number of post-ovarian inter-caecal anastomoses and lateral and medial diverticula, the median length of the terminal hook and dorsal and ventral roots of the large hamuli and egg length and width between *P. xenopodis* from the two phylogeographic lineages of the host. The WMW tests were performed and visualised via the R package *ggsignif* (Constantin & Patil, 2021).

A principal components analysis (PCA), which is commonly employed in numerical taxonomy, also that of monogeneans (e.g. Hahn *et al*., 2011), was employed to evaluate the correlation among polystomatids from different localities based upon the variance in the characters that were shown to be significantly different between the two groups. The PCA visualised whether the combination of significantly different morphological characters could discriminate between polystomatids from SA1 and SA5 hosts, despite overlap in the measurements of all these characters between the two groups, without taking into account geographical origin *a priori*. The visualisation further identified the characters that contributed most to the variation between groups. Since the Euclidean distances utilised in a PCA are sensitive to different units of measurement, the data were column-standardised beforehand in the R package *vegan* (Oksanen *et al*., 2020) as recommended by Thorpe (1981). The PCA itself was performed in base R, utilising the singular value decomposition method.

### 2.3 Molecular and phylogenetic analyses

One nuclear marker, namely *28S rDNA,* and two mitochondrial markers, namely *12S rDNA* and *COX1*, were chosen for the phylogenetic analyses. These markers have been used previously for both taxonomic and phylogenetic studies of Polystomatidae, leading to the availability of family-specific primers for these genes (Héritier *et al*., 2015, 2018; Verneau *et al*., 2009).

Extracts of DNA were obtained from six additional polystomatid specimens from six of the eight localities (Table 1) with the PCRBIO Rapid Extract PCR Kit (PCR Biosystems Ltd., London, United Kingdom). Subsequent amplification reactions were performed with 2 to 5 *μ*L extracted DNA, 1.25 *μ*L [0.2 *μ*M] forward primer and 1.25 *μ*L [0.2 *μ*M] reverse primer, 12.5 *μ*L [1 ×] master mix from the PCRBIO HS Taq Mix Red (PCR Biosystems Ltd., London, United Kingdom) and PCR grade water to the final volume of 25 *μ*L. The nuclear *28S rDNA* of the six specimens of *P. xenopodis* was amplified using the method of Verneau *et al*. (2009) with the primer pair ‘LSU5’ (5’-TAGGTCGACCCGCTGAAYTTAAGCA-3’) (Littlewood *et al*., 1997) and ‘LSU1500R’ (5’-GCTATCCTGAGGGAAACTTCG-3’) (Tkach *et al*., 1999). For the amplification of the partial mitochondrial *12S rDNA*, the thermocycling profile, forward primer ‘12SpolF1’ (5’-YVGTGMCAGCMRYCGCGGYYA-3’) and one of two reverse primers, ‘12SpolR1’ (5’-TACCRTGTTACGACTTRHCTC-3’) or ‘12SpolR9’ (5’-TCGAAGATGACGGGCGATGTG-3’), of Héritier *et al*. (2015) were used. Amplicons of the partial mitochondrial *COX1* gene were obtained with the forward primer ‘L-CO1p’ (5’-TTTTTTGGGCATCCTGAGGTTTAT-3’) and one of two reverse primers, ‘H-Cox1p2’ (5’-TAAAGAAAGAACATAATGAAAATG-3’) or ‘H-Cox1R’ (5’-AACAACAAACCAAGAATCATG-3’), also using the profile of Héritier *et al*. (2015).

For purification and sequencing, all PCR products were sent to a commercial company (Inqaba Biotec, Pretoria, South Africa) that used the ExoSAP protocol (New England Biolabs Ltd., United States) for purification and obtained the sequences with BigDye^®^ Terminator *version* 3.1 Cycle Sequencing, utilising the corresponding primer pairs used in the PCR reaction, on an ABI3500XL analyser. Sequences were assembled and manually edited in Geneious *version* 9.0 (Saint Joseph, Missouri, United States). Sequences were uploaded to GenBank (accession numbers to be added after manuscript acceptance).

The sequences from the six *P. xenopodis* specimens were aligned separately for each gene in Seaview *version* 4.7 (Gouy *et al*., 2010) with the MUSCLE algorithm *version* 3 at default settings (Edgar, 2004). For the protein-coding *COX1*, alignment was performed on the amino acid sequences, translated by the echinoderm and flatworm mitochondrial genetic code. The percentage of differing bases between the sequence pairs in each alignment was calculated in Geneious. Model-corrected pairwise genetic distances were calculated through maximum likelihood (ML) analysis in IQ-TREE *version* 2.1.2 (Minh *et al*., 2020), which first selected the optimal model of molecular evolution for each gene with the ModelFinder selection routine (Kalyaanamoorthy *et al*., 2017) with the FreeRate heterogeneity model (Soubrier *et al*., 2012) based on the Bayesian Information Criterion (BIC). The substitution models were TPM2u + F (Kimura, 1981; Soubrier *et al*., 2012) for the partial *28S rDNA*, HKY + F + I (Gu *et al*., 1995; Posada, 2003; Soubrier *et al*., 2012) for the partial *12S rDNA* and TIM2 + F + G (Posada, 2003; Soubrier *et al*., 2012; Yang, 1994) for the partial *COX1* gene. The same analyses calculated the number of invariant and parsimony informative sites for each sequence alignment.

For the subsequent phylogenetic analyses, previously published *COX1, 28S* and *12S* sequences of the closely related *P. occidentalis* (accession numbers KR856179.1, KR856121.1 and KR856160.1, respectively) were included as outgroup (Héritier *et al*., 2015) and the sequence sets were realigned as detailed above. The aligned sequences were concatenated in SequenceMatrix *version* 1.8 (Vaidya *et al*., 2011). The optimal models of molecular evolution for the *12S* and *28S rDNA* genes and the three *COX1* codon positions (Chernomor *et al*., 2016) were selected based on the BIC with the ModelFinder selection routine (Kalyaanamoorthy *et al*., 2017) implemented in W-IQ-TREE *version* 1.6.7 (Trifinopoulos *et al*., 2016). The five partitions were initially analysed separately (Chernomor *et al*., 2016) and then sequentially merged with the implementation of a greedy strategy until model fit no longer improved (Kalyaanamoorthy *et al*., 2017). The new selection procedure was implemented which included the FreeRate heterogeneity model (Soubrier *et al*., 2012). The selection routine identified three partitions in the alignment, namely the *12S rDNA* and *COX1* first codon position with best-fit model the HKY + G (Hasegawa *et al*., 1985; Yang, 1994), the *28S rDNA* and *COX1* second codon position with TIM2 + I (Gu *et al*., 1995; Posada, 2003) and the *COX1* third codon position with TIM2 (Posada, 2003).

For tree reconstruction, both ML analysis and Bayesian inference of phylogeny (BI) were performed to increase confidence in the resulting topology. The ML tree was inferred under the three partitions suggested by the selection routine. The parameter estimates were edge-unlinked for all partitions. The analysis was performed in IQ-TREE *version* 1.6.7 (Nguyen *et al*., 2015), with the assessment of branch support through ultrafast bootstrapping (UFboot2; Hoang *et al*., 2018) and the Shimodaira-Hasegawa-like (SH-like) approximate likelihood ratio test (SH-aLRT; Guindon *et al*., 2010), each with 10 000 replicates.

The BI was performed in MrBayes *version* 3.2.6 (Ronquist *et al*., 2012) implemented through the CIPRES Science Gateway *version* 3.3 on XSEDE (Miller *et al*., 2010). Posterior probabilities were calculated with four different Metropolis-coupled Markov chains over 10^6^ generations, with sampling of the Markov chain every 10^3^ generations. The first quarter of the samples was discarded as burn-in. Stationarity of the Markov chains was reached, as indicated by a deviation of split frequencies of 0.001, by a potential scale reduction factor converging to 1 and by the absence of a trend in the plot of log-probabilities as function of generations. The substitution models implemented in MrBayes were adapted from the selection of ModelFinder as the next more complex model under the BIC in terms of substitution rates available in MrBayes. Thus, the HKY model (Hasegawa *et al*., 1985) was implemented for the first partition, allowing for a discrete gamma model and the GTR (Tavaré, 1986) model was implemented with and without a proportion of invariant sites for the second and third partitions, respectively. All parameter estimates were edge-unlinked.

## 3 RESULTS

### 3.1 Morphological divergence

None of the four indicators of body size, namely body length and width and haptor length and width, were significantly different between the polystomatids from the northeastern (SA5, n = 13) and southwestern (SA1, n = 10) frog hosts (Figure 3*a*–*d*). Therefore, no adjustment was made for size in the subsequent analyses. Notably, for eight of the characters, including body length, width and haptor length, the polystomatids from southwestern hosts displayed a greater range of measurements than their counterparts from northeastern hosts (Figure 3).

**FIGURE 3.**
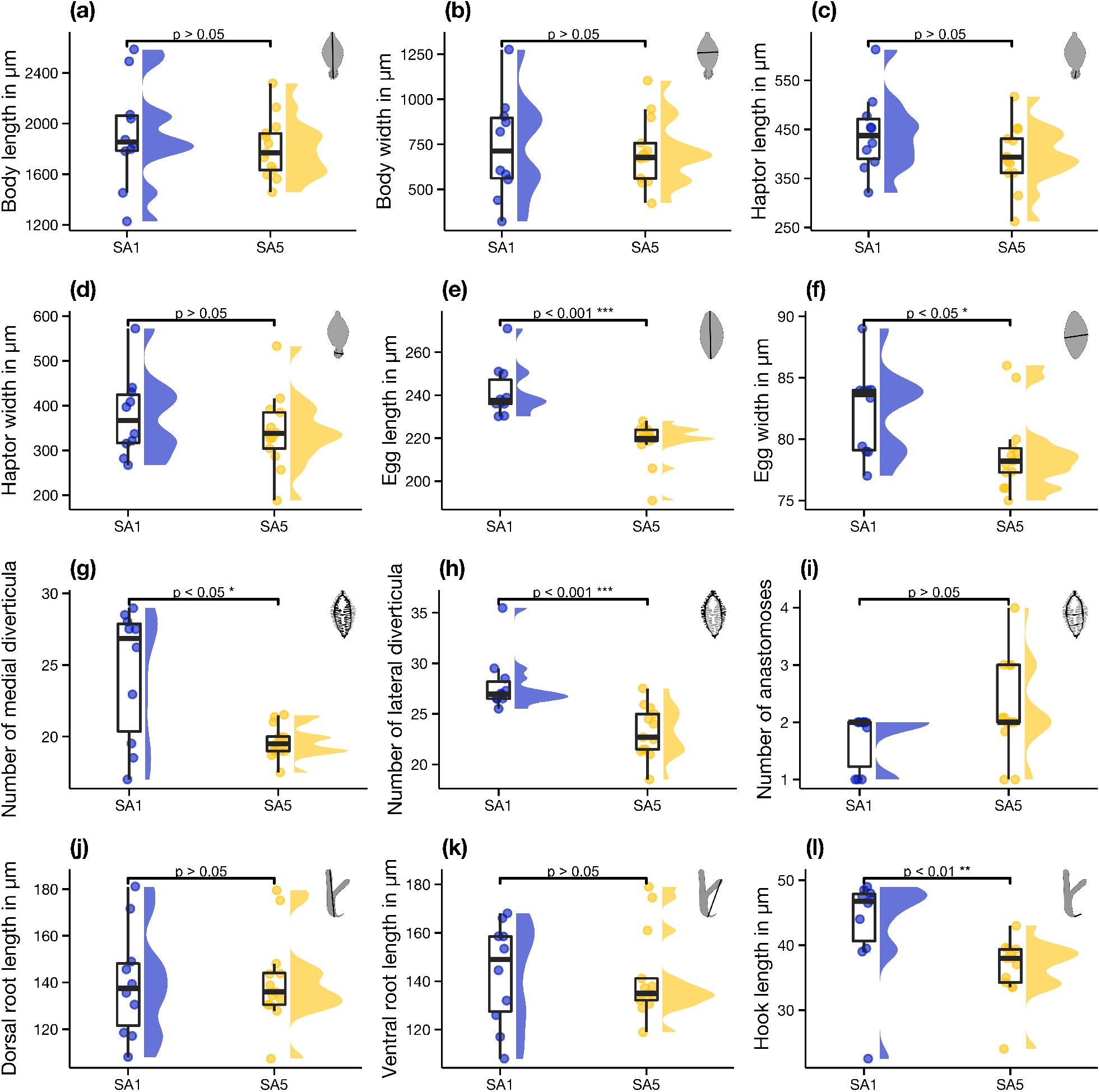
Raincloud plots of 12 morphometric characters of the parasite *Protopolystoma xenopodis*, compared based on the geographic origin of their host *Xenopus laevis*, namely SA1 and SA5. The characters are (a) body length, (b) width, (c) haptor length, (d) width, (e) egg length, (f) width, (g) number of medial, (h) lateral diverticula and (i) post-ovarian intercaecal anastomoses, (j) length of the dorsal root, (k) ventral root and (l) terminal hook of the large hamuli. Points indicate raw data along with their distributions to the right and summary statistics, the first, second and third quartiles, are given in boxplots to the left. The brackets above the plots indicate the significance levels calculated by Wilcoxon-Mann-Whitney tests that compared the characters between the two groups.

Polystomatids from southwestern frog hosts had significantly longer and wider eggs than those from northeastern hosts (Figure 3*e*–*f*). There were marked differences in the gut morphology between the polystomatids from the two regions. Polystomatids from southwestern frog hosts had significantly more medial and lateral diverticula of the caeca than those from the northeastern hosts (Figure 3*g*–*h*). On the other hand, even though there were some northeastern polystomatids with up to four post-ovarian intercaecal anastomoses, as opposed to their southwestern counterparts where no specimen had more than two, there was no significant difference in this character between polystomatids from these two regions (Figure 3*i*). In terms of large hamulus shape and size, there was no overall difference in the length of the dorsal and ventral roots of the large hamuli between polystomatids from the two regions (Figure 3*j*–*k*). However, southwestern polystomatids had significantly longer terminal hooks than the northeastern polystomatids (Figure 3*l*).

Thus, *P. xenopodis* from the southwestern region displayed less variation in the number of anastomoses (SA1_min:max_ = 1–2; SA5_min:max_ = 1–4), possessed more diverticula, both laterally (SA1_median_ = 27; SA5_median_ = 23) and medcially (SA1_median_ = 28; SA5_median_ = ^20)^, had longer terminal hooks of the hamuli (SA1_median_ = 46.75 *μ*m; SA5_median_ = 36.50 *μ*m) and had longer (SA1_median_ = 250.0 *μ*m; SA5_median_ = 220.5 *μ*m) and wider eggs (SA1_median_ = 84 *μ*m; SA5_median_ = 78 *μ*m) than their northeastern counterparts. Moreover, egg length was a diagnostic character, with no overlap in measurements observed between the two groups of polystomatids (SA1_min:max_ = 236–271 *μ*m; SA5_min:max_ = 191–228 *μ*m).

According to the results of the PCA, the combination of the five significantly different morphological characters, namely egg length and width, terminal hook length and number of lateral and medial diverticula, allowed reasonable discrimination between the polystomatids from the two host lineages (Figure 4). The first two principal components (PCs) accounted for 64.54% and 14.04% of the observed variance, together explaining 78.58% of the variance in the data. The loadings of PC1 and PC2 were both positive and negative.

**FIGURE 4.**
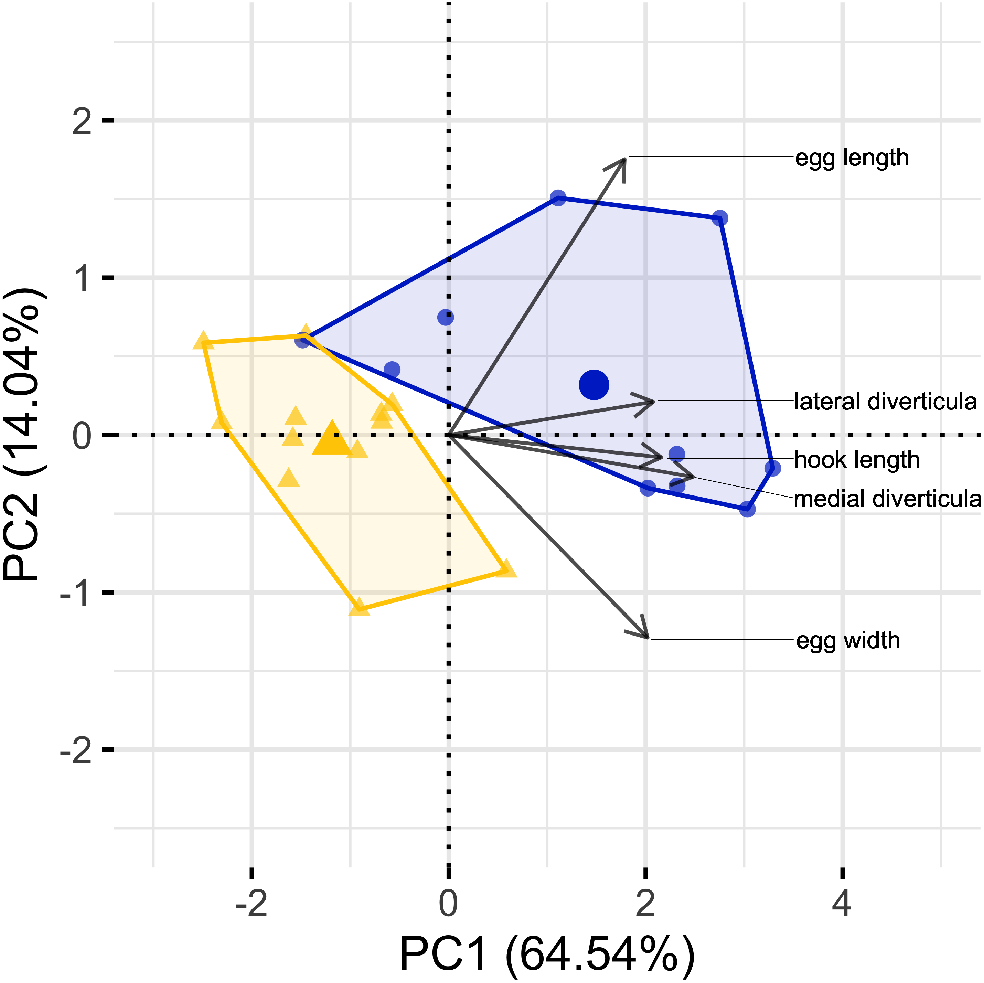
The first two principal components derived from five morphometric variables—length and width of the egg, length of the terminal hooks of the hamuli (in *μ*m) and the number of lateral and medial diverticula—of 23 *Protopolystoma xenopodis* associated with the southwestern (SA1, circles in blue) and northeastern (SA5, triangles in yellow) phylogeographic lineages of its frog host *Xenopus laevis*.

### 3.2 Phylogenetic divergence

In the case of the partial *28S rDNA* alignment without the outgroup sequence, a total of 1721 bases contained 18 variable and 7 parsimony informative sites. Model-corrected genetic distances in the *28S rDNA* sequences among the six specimens ranged from 0 to 1.6% (Table 2). The partial *12S rDNA* data set without the outgroup sequence was represented by an alignment of 505 base pairs. Of the 505 sites, 67 sites were variable and 23 were parsimony informative. Model-corrected genetic distances in the *12S rDNA* sequences among the six *P. xenopodis* specimens ranged from 0.12 to 3.77% (Table 2). For the *COX1* gene alignment, the data set without the outgroup amounted to 418 base pairs, where 25 of the 74 variable sites were parsimony informative. Model-corrected genetic distances in the *COX1* sequences among the six specimens ranged from 0.14 to 9.00% (Table 2). The concatenation of the three alignments with the outgroup sequences, which was used for the subsequent phylogenetic analyses, yielded a total of 2667 base pairs. There were 250 variable sites, of which 93 were parsimony informative.

**TABLE 2.** Pairwise genetic distances (%) of three partial gene sequences from six specimens of *Protopolystoma xenopodis* from *Xenopus laevis* collected at six localities, here named according to the nearest town. Model-corrected distances are given above the diagonal and percentage of non-identical bases are given below.

|  | CPT | HRM | MDM | DLS | TZN | PTC |
| --- | --- | --- | --- | --- | --- | --- |
| **Nuclear 28S rDNA** |  |  |  |  |  |  |
| Cape Town (CPT) | - | 0.05 | 0.42 | 0.40 | 0.50 | 0.56 |
| Hermanus (HRM) | 0.09 | - | 0.37 | 0.36 | 0.61 | 0.48 |
| Modimolle (MDM) | 0.53 | 0.59 | - | 0.74 | 1.60 | 0.85 |
| Dullstroom (DLS) | 0.47 | 0.53 | 0.88 | - | 0.00 | 0.07 |
| Tzaneen (TZN) | 0.47 | 0.53 | 0.88 | 0.00 | - | 0.05 |
| Potchefstroom (PTS) | 0.59 | 0.65 | 0.10 | 0.12 | 0.06 | - |
| **Mitochondrial 12S rDNA** |  |  |  |  |  |  |
| Cape Town (CPT) | - | 2.17 | 2.02 | 2.15 | 2.15 | 2.11 |
| Hermanus (HRM) | 9.76 | - | 1.39 | 3.77 | 1.37 | 1.37 |
| Modimolle (MDM) | 10.18 | 4.50 | - | 2.45 | 0.41 | 0.40 |
| Dullstroom (DLS) | 10.14 | 6.52 | 6.94 | - | 2.87 | 2.70 |
| Tzaneen (TZN) | 9.96 | 4.08 | 1.02 | 6.52 | - | 0.12 |
| Potchefstroom (PTS) | 10.37 | 4.08 | 1.02 | 6.92 | 0.41 | - |
| **Mitochondrial COX1 gene** |  |  |  |  |  |  |
| Cape Town (CPT) | - | 9.00 | 9.00 | 9.00 | 9.00 | 9.00 |
| Hermanus (HRM) | 11.99 | - | 0.24 | 9.00 | 0.29 | 0.32 |
| Modimolle (MDM) | 12.98 | 4.09 | - | 9.00 | 0.20 | 0.23 |
| Dullstroom (DLS) | 12.23 | 7.43 | 8.89 | - | 9.00 | 9.00 |
| Tzaneen (TZN) | 13.19 | 3.84 | 2.40 | 9.11 | - | 0.14 |
| Potchefstroom (PTS) | 12.47 | 4.32 | 2.64 | 8.39 | 1.68 | - |

In agreement with the morphometric analyses, phylogenetic tree reconstruction based on the concatenated *12S* and *28S rDNA* and *COX1* gene alignments revealed remarkable divergence in *P. xenopodis* based upon geographic origin. Polystomatids from the SA5 localities formed a well-supported clade (Figure 5). *Protopolystoma xenopodis* from Hermanus and Cape Town (SA1) were earlier diverging than those from the SA5 localities and were rendered paraphyletic by the SA5 lineage in both the BI and ML analyses. Additionally, the BI could not resolve the relationships between *P. xenopodis* from Dullstroom, Tzaneen and Potchefstroom (SA5), even though the sister relationship of *P. xenopodis* from Potchefstroom and Tzaneen to *P. xenopodis* from Dullstroom had high support in the ML.

**FIGURE 5.**
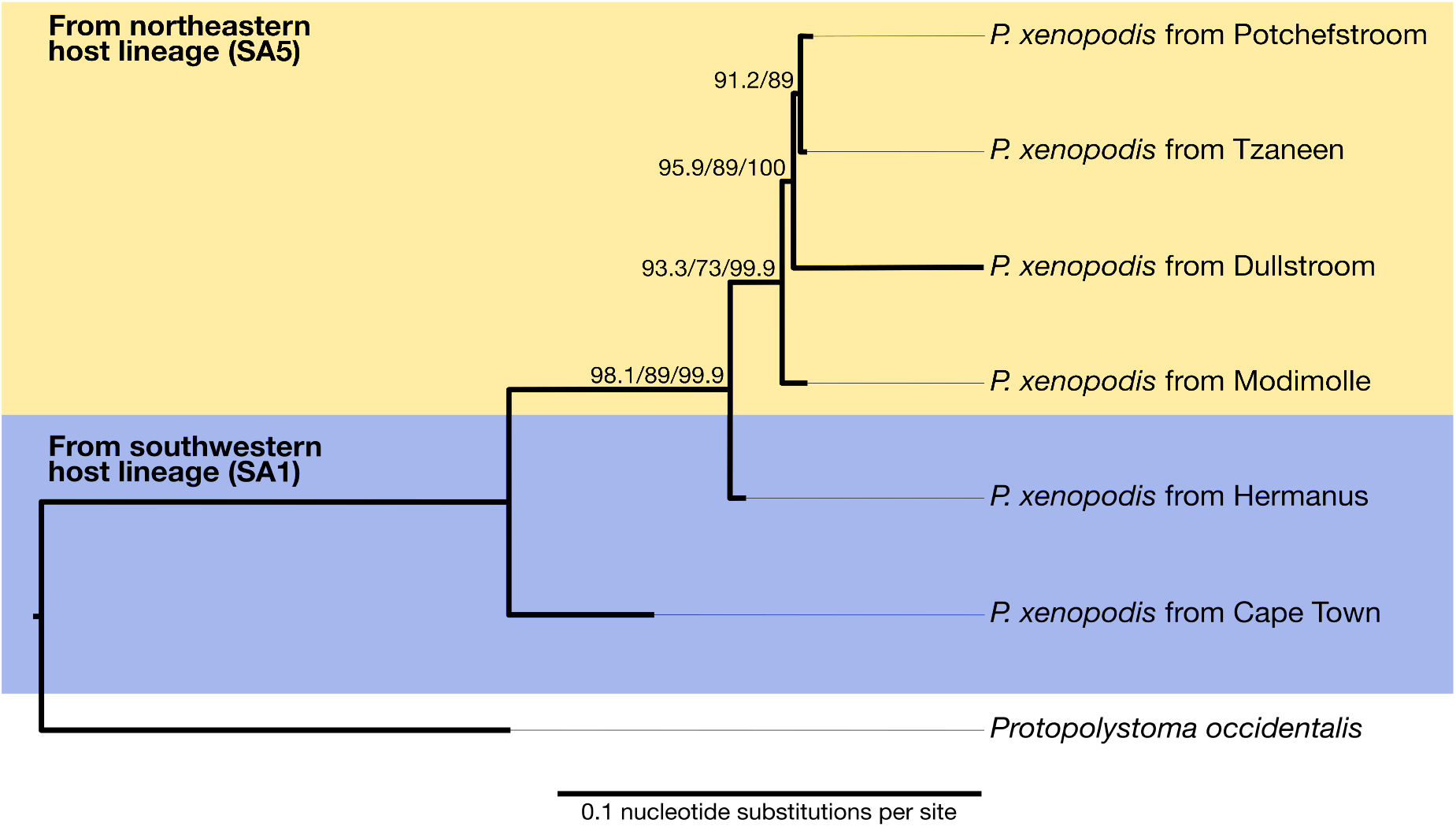
Maximum likelihood consensus phylogram of six *Protopolystoma xenopodis* specimens, inferred from the *COX1* gene and *12S* and *28S rDNA* sequences. The polystomatids were recovered at six localities, indicated by the name of the nearest town, from *Xenopus laevis* frog hosts from two phylogeographic lineages (SA1 in blue, SA5 in yellow). The closely related *P. occidentalis* from *X. muelleri* from Togo was used to root the phylogram. Values at the nodes indicate support, where available, as calculated by ultrafast bootstrapping (first value), SH-like approximate likelihood ratio test (second value) and posterior probabilities (third value).

## 4 DISCUSSION

The present investigation is the first integrative approach to the intraspecific diversity of *P. xenopodis*, the widespread bladder parasite of the globally invasive frog *X. laevis*. Both morphological and molecular data reveal notable intraspecific divergence in *P. xenopodis* collected from two lineages of their host *X. laevis* in South Africa. The combination of egg length and width, number of diverticula of the gut and length of the terminal hook of the large hamulus provides a set of key characters that differentiate northeastern and southwestern populations of *P. xenopodis* in South Africa. Furthermore, the morphological differentiation is supported by the results of the multilocus phylogenetic analyses. Moreover, this intraspecific divergence corresponds to the documented phylogeographic structuring of the host *X. laevis* in its native range (Furman *et al*., 2015).

Intraspecific variation in morphological characters has been reported before in many species of Polystomatidae and herein *P. xenopodis* is no exception. Especially the number of inter-caecal anastomoses and medial and lateral diverticula are suggested as highly variable characters in polystomatid monogeneans, including *P. xenopodis* (e.g. Aisien & du Preez, 2009; du Preez *et al*., 2002; Tinsley, 1974, 1978; Tinsley & Jackson, 1998b). Likewise, the high variability in the length of the terminal hook of the large hamulus and genital spine length in *P. xenopodis* has been pointed out previously (Tinsley & Jackson, 1998b). Yet, genital spine length in *P. xenopodis* is the only character for which the link between morphological variation and geographic distance has been explored to date (Tinsley & Jackson, 1998b). Unfortunately, genital spine length could not be measured in the present study. The mounting procedure that we applied to the available specimens, whilst it is ideal for measurements of the soft structures and large hamuli, does not allow for sufficient flattening of the specimens to ensure that the smaller sclerites, such as the genital spines and marginal hooklets, are mounted horizontally.

In accordance with the variation in morphometrical characters, the mitochondrial *COX1* gene and *12S rDNA* show remarkable intraspecific divergence both within and between the southwestern and northeastern clusters of *P. xenopodis*. In fact, the divergence in the *COX1* gene of *P. xenopodis* far exceeds that of *Madapolystoma* spec. from frog hosts across Madagascar (between 1.7 and 13.2% for *P. xenopodis* and 0.3 and 1.8% in *Madapolystoma* spec.), the only other polystomatid genus for which intraspecific genetic variation has been assessed to date (Berthier *et al*., 2014). This points towards the nuclear *28S rDNA* as a more useful marker for species recognition in *Protopolystoma*. Yet, even for the *28S rDNA*, intraspecific divergence in *P. xenopodis* is generally higher than what was reported for *Madapolystoma* spec. (between 0.1 and 1.6% for *P. xenopodis* and 0.08 and 0.23% in *Madapolystoma* spec.) (Berthier *et al*., 2014). The observed geographic variation suggests that future taxonomic studies of Polystomatidae should make use of geographically representative data sets, both when relying on traditional morphometric and advanced molecular approaches.

Nonetheless, given the overlap in measurements between the specimens of *P. xenopodis* from different geographic regions, we assume that the observed divergence is still within the bounds of intraspecific variation. It is important to keep in mind that the examined parasites hail from the two most geographically distanced lineages of the host (de Busschere *et al*., 2016; Furman *et al*., 2015). With the addition of specimens collected from the frog hosts that hail from localities between these two areas, geographically speaking, it is not unrealistic to imagine that morphological variation will present itself on a spectrum that correlates with geographical distance. This is in accordance with the “significant, but continuous” variation in genital spine length that was observed in the study by Tinsley & Jackson (1998b). Likewise, similar investigations of fishes and reptiles revealed that potentially interesting phenotypic divergence was initially reported simply because only the extremes of the distributional range were considered—once the intermediate populations were included in the analyses, phenotypic variation represented geographical variation along a cline (e.g., Ennen *et al*., 2014; Manier, 2004; Risch & Snoeks, 2008; van Steenberge *et al*., 2011, 2015).

On the face of it, one could interpret the observed morphometric variation against the backdrop of the different climatic conditions on either side of the Great Escarpment where our specimens were collected, namely summer rainfall regime to the northeast and a winter rainfall regime to the southwest. Indeed, phenotypic plasticity is a common response to changes in environmental conditions in representatives of Monogenea. This is true of the shape of the hamuli in gyrodactylid species, which has been shown to vary with temperature in isogenic lineages (Olstad *et al*., 2009). Moreover, both Harris (1998) and Hahn *et al*. (2011) suggested that most of the infrapopulation morphological variation in gyrodactylids can be ascribed to environmental drivers, since it could not be reliably linked to genetic differences. In the case of a capsalid monogenean, temperature differences under experimental conditions drove differences relating to body size, but not relating to the size and shape of sclerotised features (Brazenor *et al*., 2018). In *P. xenopodis*, temperature can influence egg production rate in the laboratory (Jackson & Tinsley, 1988). Lamentably, the influence of temperature on egg dimensions has not been investigated. All in all, there is evidence that environmental parameters, especially temperature fluctuations, can drive morphological variation in Monogenea within populations or under experimental conditions. However, it is less likely to be the cause of between-population morphological variation, as is revealed by our study.

When both the morphological and molecular lines of evidence are considered, it becomes clear that the observed variation in *P. xenopodis* is not merely the product of plasticity during ontogenic development in response to differing climatic conditions. Firstly, the morphological differentiation between the specimens from the two host lineages involves some of the characters that are important for species delineation in the genus, such as gut and hamulus morphology (Tinsley & Jackson, 1998b). This hints at a link with incipient speciation. Secondly, the observed morphological differences correspond to marked phylogenetic divergence on the intraspecific level in both mitochondrial and nuclear genes of *P. xenopodis*. This divergence echoes the phylogeographic structuring of its frog host across South Africa (Furman *et al*., 2015), hinting at congruence between the intraspecific diversification of *X. laevis* and *P. xenopodis*, which may be explored in future studies.

The Great Escarpment is a well-studied landscape barrier that seems to have shaped the diversification or restricted the expansion of a great many species in South Africa, including representatives of insects, frogs, snakes, lizards and small mammals (e.g. Barlow *et al*., 2013; Makokha *et al*., 2007; Mynhardt *et al*., 2015; Nielsen *et al*., 2018; Predel *et al*., 2012). This geological feature likely also had an impact on population structure within the African Clawed Frog *X. laevis* (Furman *et al*., 2015). Thus, in the light of the high host specificity and ancient association of *Xenopus* and *Protopolystoma* species (Tinsley & Jackson, 1998a), it comes as no surprise that the phylogeographic divergence in *X. laevis* on either side of the Escarpment is mirrored in the morphological, and especially phylogenetic, divergence of *P. xenopodis*. Nonetheless, the Escarpment is no barrier to the well-documented human-mediated domestic translocation of *X. laevis* from the southernmost part of its range to other localities in southern Africa (Measey & Davies, 2011; van Sittert & Measey, 2016). This widespread phenomenon could also contribute to the spread of co-translocated southernmost *P. xenopodis* to the northern parts of its range, which is clearly possible when one considers *P. xenopodis*’ co-intruduction into the invasive range (Schoeman *et al*., 2019). As a next step, more detailed investigations that consider the phylogeography of corresponding host-parasite pairs could shed light on the evolutionary and ecological repercussions of the anthropogenic movement of *X. laevis* in southern Africa.

In sum, there are clear indications of geographic variation in *P. xenopodis* in South Africa, despite the low sample sizes and patchy geographic presentation of the present study. The findings of this exploratory study open new avenues of investigation for this widespread host-parasite system. Based upon the integration of morphometry and multilocus phylogenetics, our findings bring to light a possible link between the evolutionary histories of both frog host and flatworm parasite. The corresponding morphological and molecular divergence of both *X. laevis* and *P. xenopodis* is a factor to keep in mind in terms of their ability to colonise and adapt to new environments, as was noted for invasive *X. laevis* in France (de Busschere *et al*., 2016). In addition, the phylogeographic analysis of *P. xenopodis* has the potential to act as a key piece of evidence in the reconstruction of the invasion histories of *X. laevis,* as has been demonstrated in a handful of other studies on the monogenean parasites of invasive fish (Huyse *et al*., 2015; Kmentova *et al*., 2019; Ondračková *et al*., 2012). Ultimately, the newly revealed geographic variation in the most common parasite of *X. laevis* demonstrates that we have barely scratched the surface when it comes to understanding the native parasite dynamics of the world’s most widespread amphibian.

## ACKNOWLEDGEMENTS

The authors express their sincere thanks to the farm and smallholding owners who graciously gave permission for collection to take place on their properties and who provided lodging for the research team: Fanus and Olga Kritzinger, Tobie Bielt and Gert Bench. In addition, we thank Mathys Schoeman and Roxanne Viviers who also assisted with fieldwork. The utilisation of the frogs and the research protocols were approved by the Animal Care, Health and Safety in Research Ethics (AnimCare) Committee of the Faculty of Health Sciences of the North-West University (ethics number: NWU-0380-16-A5-01). Animals were sampled under the permit 0056-AAA007-00224 (CapeNature) provided by the Department of Economic Development, Environmental Affairs and Tourism. Special Research Funds (BOF) of UHasselt supported MPMV (no. BOF20TT06) and NK (no. BOF21PD01). We further acknowledge the financial support of the National Research Foundation (NRF) of South Africa. ALS received funding from the DSI-NRF Centre of Excellence for Invasion Biology and from the NRF South African Institute for Aquatic Biodiversity. LHdP is indebted to the NRF Foundational Biodiversity Information Programme (no. 120782) for financial support. Any opinion, findings and conclusions or recommendations expressed in this material are those of the authors and the NRF does not accept any liability in this regard.

